# Individualized Cerebellar Damage Predicts the Behavioral Disorders in Children with Brainstem Tumors

**DOI:** 10.1101/2024.08.01.606261

**Authors:** Heyuan Jia, Kaikai Wang, Mingxin Zhang, Guocan Gu, Yiying Mai, Xia Wu, Congying Chu, Xuntao Yin, Peng Zhang, Lingzhong Fan, Liwei Zhang

**Affiliations:** School of Instrumentation and Optoelectronic Engineering, Beihang University, Beijing, China; Institute of Large-scale Scientific Facility and Centre for Zero Magnetic Field Science, Beihang University; Brainnetome Center, Institute of Automation, Chinese Academy of Sciences, Beijing 100190, China; Department of Neurosurgery, Beijing Tiantan Hospital, Capital Medical University, Beijing, China; Department of Radiology, Guangzhou Women and Children’s Medical Center, Guangzhou Medical University, Guangzhou, China; School of Health and Life Sciences, University of Health and Rehabilitation Sciences, Qingdao 266000, China; China National Clinical Research Center for Neurological Diseases, Beijing Tiantan Hospital, Capital Medical University, Beijing, China; Beijing Neurosurgical Institute, Beijing Tiantan Hospital, Capital Medical University, Beijing 100070, China

**Author notes:** Correspondence to: Liwei Zhang, Professor, Department of Neurosurgery, Beijing Tiantan Hospital, Capital Medical University, Beijing, China, Lingzhong fan, Professor, Brainnetome Center, Institute of Automation, Chinese Academy of Sciences, Beijing 100190, China, Zhang Peng, M.D., Department of Neurosurgery, Beijing Tiantan Hospital, Capital Medical University, Beijing, China. These authors contributed equally to this work (Heyuan Jia, Kaikai Wang).

**Keywords:** Brainstem tumor, cerebellar atrophy, social problem, normative model, machine learning

## Abstract

**Background and Purpose:** Brainstem tumors are rare but cause enduring behavioral issues, challenging patients and surgeons. Research on cerebellar changes in these patients limited, despite symptoms similar to cerebellar injuries. This study aims to investigate cerebellar damage pattern resulting from brainstem tumors and its association with behavioral disorders.

**Methods:** In this study, a U-Net-based segmentation algorithm was used to divide the cerebellum into 26 lobules, which were then used to build a normative model for assessing individual structural deviations. Furthermore, a behavior prediction model was developed using the total outlier count (tOC) index and brain volume as predictive features.

**Results:** Most patients were found to have negative deviations in cerebellar regions, particularly in anterior lobules like Left V. Higher tOC was significantly associated with severe social problems (r = 0.31, p = 0.001) and withdrawal behavior (r = 0.28, p = 0.001). Smaller size of cerebellar regions strongly correlated with more pronounced social problems (r = 0.27, p = 0.007) and withdrawal behavior (r = 0.25, p = 0.015). Notably, lobules Right X, V, IV, VIIB, Left IX, VIII, and X influenced social problems, while Left V, Right IV, Vermis VI, and VIII impacted withdrawal behavior.

**Conclusions:** Our study revealed cerebellar damage patterns in patients with brainstem tumors, emphasizing the role of both anterior and posterior cerebellar lobes in social problems and withdrawal behavior. This research sheds light on the brain mechanisms underlying complex behavioral disorders in brainstem tumor patients.

## Introduction

Brainstem tumors originate specifically in the medulla oblongata, pons, and midbrain. They constitute 15% of pediatric brain tumors, with diffuse intrinsic pontine gliomas (DIPG) making up 80% of these cases^1^. Unfortunately, DIPG in children is associated with a very poor prognosis and leads to complex neurological dysfunction. Historically, higher cognitive functions were not attributed to the brainstem^2^. However, recent reports have indicated abnormal cognitive and behavioral disorders in patients with brainstem lesions, often masked by severe neurological symptoms^3^. Notably, symptoms like behavioral disorders, withdrawal behaviors, attention deficits, social disturbances, irritability, and flattened affect are gaining clinical attention. These symptoms can persist long after surgical intervention^4^. Recent neuropsychological research has demonstrated widespread behavioral impairments in children with brainstem tumors^5^, aligns with previous reports on symptoms caused by posterior fossa tumors^6,7^. Researchers believe these behaviors may result mainly from cerebellar damage^8^. However, systematic investigations into indirect cerebellar changes and their relationship with cognitive functions in brainstem tumor patients are limited.

The cerebellum, located inferiorly and posteriorly to the cerebral cortex, overlays the pons, medulla and its intimately connected to the brainstem via dense neural fiber bundles^9^. The corticospinal tract and spinocerebellar tract connect cortical brain regions with the contralateral cerebellum through the intermediary pontine region^10^. Additionally, fascicular pontocerebellar fibers extend extensively into the opposite hemisphere, forming crucial cerebellar peduncles^11^. Moreover, the cerebellum also connects with the hypothalamus, limbic system, and various brainstem regions such as the ventral tegmental area^12^. The close spatial relationship predisposes the growth of brainstem tumors to potentially the normal functioning of the cerebellum^13^. Current research suggests that the brainstem engages in cognitive processing through its interaction with the frontal lobe system and also plays a role in cognitive processing as an integral part of the cerebro-cerebellar circuit^14^. Clinical observations support the hypothesis, showing that ipsilateral cerebellar atrophy is common in brainstem tumor cases, as well as in brainstem ischemia, stroke, and trauma^15^. This may be due to damage to the direct fiber connections between the brainstem and cerebellum^16^.

In recent years, research has primarily focused on the non-motor functions of the cerebellum. The anterior regions of the cerebellum, including lobules I-IV and lobule VIII, are mainly associated with motor functions, while the posterior and lateral regions are closely linked to non-motor functions^17^. Functional imaging studies have shown that various cognitive tasks can activate the posterior cerebellum. For instance, during social cognitive tasks, regions such as Crus I and II in the posterior cerebellum are significantly activated, indicating its crucial role in social understanding and learning sequences of social behavior^18^. Schmahmann identified the cerebellar cognitive affective syndrome (CCAS), resulting from posterior cerebellar damage. It is characterized by executive dysfunction, spatial cognitive impairments, language disorders, and changes in personality and emotion^19^. Notably, the impairments caused by these damages are similar to the symptoms caused by brainstem tumors^20^. Thus, we hypothesize that the complex behavioral disorders observed in patients are caused by heterogeneous alterations in the cerebellum resulting from brainstem tumors.

To test our hypothesis, we employed the ACAPULCO (Automatic Cerebellum Anatomical Parcellation using U-Net with Locally Constrained Optimization) for effective cerebellar segmentation of deformed cerebellum. Additionally, we established a normative model to quantify cerebellar damage pattern by integrating data from healthy children. This model allowed us to assess individual deviations in brainstem tumor patients. Subsequently, we used machine learning to link these deviations to specific behaviors, identifying cerebellar regions associated with each behavior. Through these investigations, we sought to uncover the underlying brain mechanisms responsible for behavioral impairments in patients with brainstem tumors.

## Methods

### Participants

A cohort of 171 pediatric patients diagnosed with brainstem tumors at the Department of Neurosurgery, Beijing Tiantan Hospital, was enrolled between April 2019 and December 2022. Inclusion criteria were as follows: 1) Initial diagnosis of brainstem tumor with no prior treatment; 2) Tumors confined to the brainstem without infiltrating the cerebellum, spinal cord, or thalamus; 3) Absence of comorbid psycho-neurological disorders, behavioral and personality disorders; 4) No history of developmental disorders. All patients underwent 3DT1 magnetic resonance imaging (MRI) during their initial visit and completed the Child Behavior Checklist (CBCL) scale - Parent version. Informed consent was obtained as per the protocol approved by the Beijing Tiantan Hospital, Capital Medical University, in accordance with the 1964 Declaration of Helsinki and its later amendments.

To establish a normative model, data from 849 healthy children were collected from publicly available databases, including 61 locally recruited in Beijing, and 69 from the fcon_1000.projects dataset, 450 from the Child Connectome Project (CCNP), and 269 from the Children’s Hospital in Guangzhou. After excluding subjects with severe cerebellar compression or poor image quality, our final analysis included 147 patients and 849 healthy children. Fig. 1 illustrates the enrollment process in a flowchart. We downloaded MRI T1 images and corresponding phenotypic information from these databases, with the demographic characteristics of the subjects summarized in Table 1.

**Figure 1.**
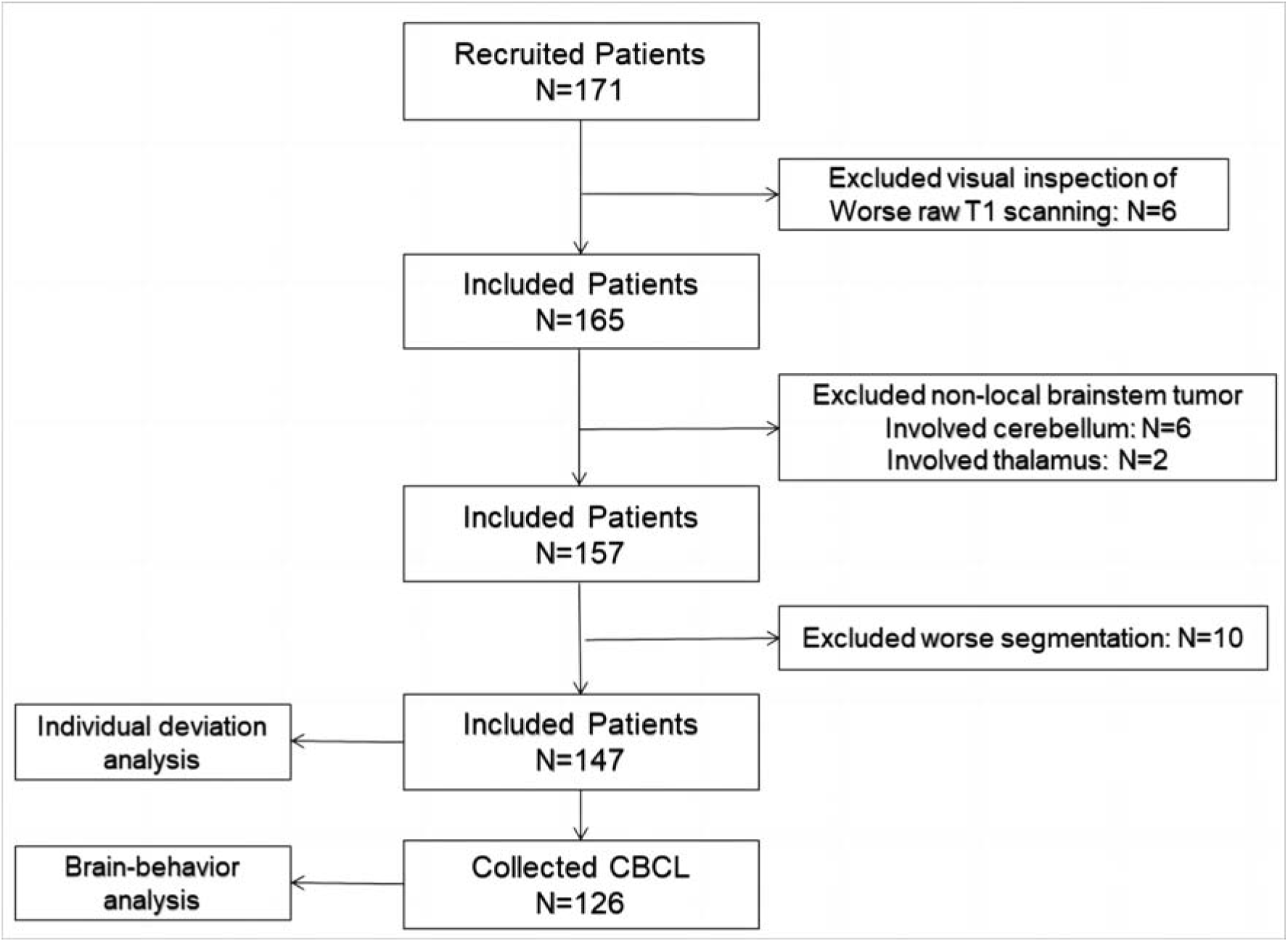
The flow chart of patient enrollment.

**Table 1.**
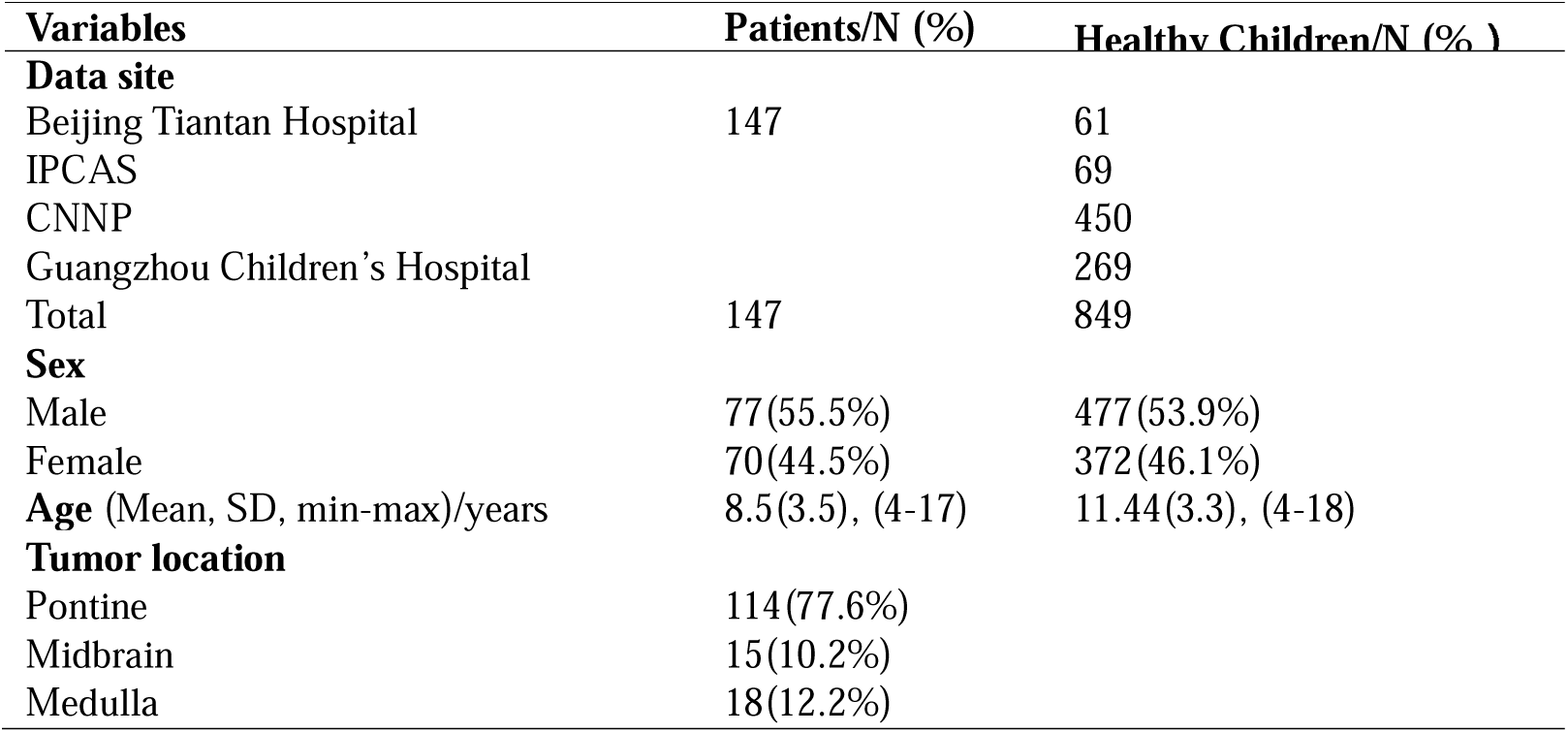
The demographic of cohort and clinical characteristic.

### MRI acquisition and processing

Structural MRI scans were performed using a 3.0 T scanner (Ingenia CX, Philips Healthcare) with a 32-channel head receiver coil. The 3D T1-weighted sequence consisted of 196 contiguous sagittal slices with a voxel size of 1.0 x 1.0 x 1.0 mm³. Specific parameters were: TR = 6.572 ms; TE = 3.025 ms; FA = 8°; slice thickness = 1 mm; in-plane resolution = 1×1 mm. DICOM images were converted to NIFTI format. The typical radiological characteristics of patients with brainstem tumors are presented in Fig. 2. To ensure accurate segmentation and avoid misclassifying the brainstem as the cerebellum, neurosurgeons delineated the brainstem as the volume of interest (VOI) using MRIcroGL. Automatic segmentation of cerebellar lobules was performed using ACAPULCO software (version 0.2.1) with a pediatric template^21^. Segmentation accuracy was visually inspected by two researchers, with manual corrections made using ITK-SNAP. We obtained volumes for 28 distinct cerebellar regions, including left and right lobules I-III, IV, V, VI, VIIB, VIIIA, VIIIB, IX, X, Crus I, Crus II, and vermis lobules VI, VII, VIII, IX, X, and corpus medullare. To address the non-uniform margin between lobules VIIIA and VIIIB, they were combined into a single lobule, designated as lobule VIII. Thus, 26 independent cerebellar region volumes were used to analyze individualized cerebellar changes and their associations with behavioral outcomes.

**Figure 2.**
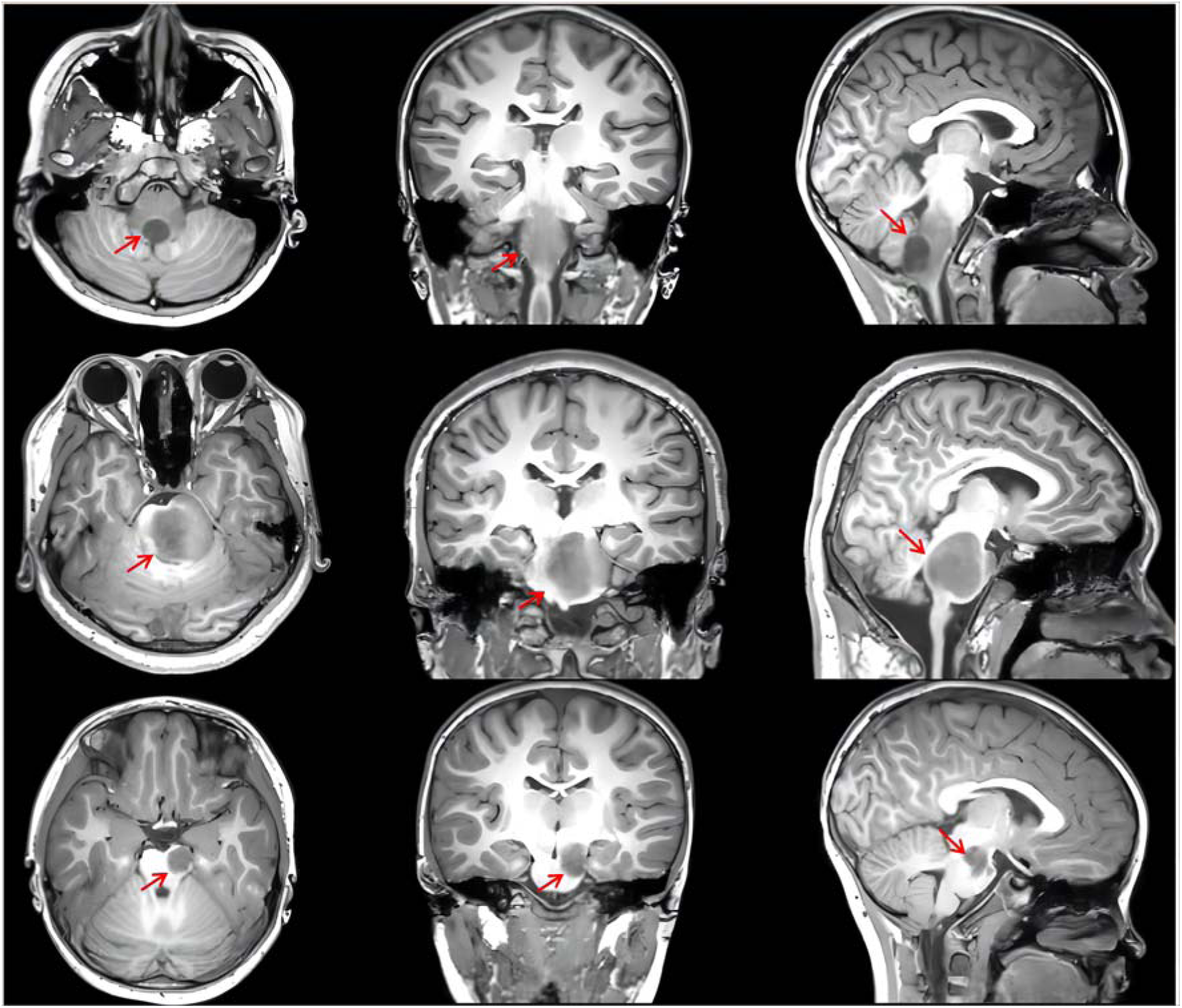
The radiological characteristics of patients with brainstem tumors. **(A-C)** The 3D-T1 weighted MRI images from left to right, representing axial, coronal, and sagittal positions, respectively. Moving from top to bottom, the primary tumor is located in the medulla oblongata, pons, and midbrain. The red arrow highlights the tumor tissue, characterized by an uneven low signal.

### Behavioral assessment

Participants’ behavioral profiles were assessed using the Achenbach Child Behavioral Checklist (CBCL), comprising 113 items categorizing behavioral issues into eight factors: somatic symptoms, withdrawal behaviors, depression/anxiety, aggressive behavior, delinquent behavior, thought problems, and social problems^22^.

### Data harmonization

We utilized the Combat method to harmonize data from different sources, mitigating non-biological variance from inter-scanner discrepancies. This method has been validated for effectively reducing unwanted variation introduced by site differences in structure MRI investigations^23^. Notably, this method retained age, gender, and TIV as covariates while excluding diagnostic labels.

### Normative modeling

Using volumetric data from the 26 cerebellar regions, we established normative models with Gaussian Process Regression (GPR), incorporating age and gender as covariates^24^. This Bayesian non-parametric modeling approach inherently considers prediction uncertainty and provides probability distributions for predicted values. We initially estimated models with 10-fold cross-validation on the healthy control dataset, then developed definitive models for deviation analysis in patients. For each participant, the normative model was used to predict volume estimates for each region, along with corresponding predictive uncertainty. Predictive uncertainty contours were then used to characterize centile variations within the cohort, enabling us to parameterize volume deviations relative to the reference cohort.

### Estimation of cerebellar volume deviations for each subject

For each glioma patient, we utilized normative percentile charts from healthy control to assess individual deviations in cerebellar region volumes. Deviations were quantified by computing a Z-score based on observed and predicted volume values for each region. For each patient i, the Z score of a cerebellum region j was calculated using the formula:

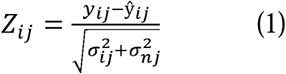

where *y_ij_* is the true volume value, *ŷ_ij_* is the expected volume value estimated from the GPR, *σ_ij_* is the predictive uncertainty, and *σ_nj_* is the variance learned from the normative distribution n.

### Statistical Analysis

#### Deviation rate calculation and statistics

Using the normative model, the relative deviation distribution for each patient was compared to the general norm. The deviation rate was computed by counting patients with Z-values exceeding ±1.96 and dividing by the total patient count. This yielded positive and negative deviation rates, indicating the extent of deviations in each brain region.

#### Outlier definition

Outliers indicating diminished cerebellar volume were identified within each region, precisely defined as Z < - 1.96, corresponding to the lower 2.5th percentile of the normative cerebellar volume distribution. We selected the lower threshold for outlier detection to focus on evaluating reductions in cerebellar volume associated with glioma. The total outlier count (tOC) across all 26 regions was computed for each participant^25^.

#### tOC associations with social problems and withdrawal behaviors

We investigated the relationship between tOC and clinical symptoms, specifically social problems and withdrawal behaviors, using Pearson correlation analysis. To determine if tOC could indicate essential behaviors in patients, we employed a ridge regression model with a nested fivefold cross-validation (5F-CV). The outer loop assessed the model’s generalization to unseen subjects, and the inner loop performed a grid search to identify the optimal parameter within a specified A range ([2^−10^, 2^−9^, … , 2^−4^, 2^−5^]. We conducted 101 repetitions of the 5F-CV, calculating prediction accuracy as the mean r value of the five folds, resulting in a final accuracy of the median r value from 101 replicates. Linear regression was applied within each fold to adjust for age and sex effects. Permutation testing assessed the prediction’s significance by estimating the empirical distribution of prediction accuracy under the null hypothesis of no association between cerebellar volume deviations and behavior values. The prediction procedure was repeated 1000 times to create the null distribution. In each iteration, behavior scores were permuted across the training samples^26^.

#### Prediction of behaviors from cerebellar volume deviations using ridge regression

After analyzing the relationship between tOC and behavior, we investigated whether cerebellar volume deviations captured significant behavioral characteristics in patients. We combined the Z scores from 26 regions into a comprehensive feature vector representing an individual’s deviation, and used these to adjust the ridge regression model. Each region was assigned a feature weight indicating its importance and negative or positive association with behavior. We evaluated feature weights across 101 iterations, focusing on the median r value run. The significance of each feature weight was determined through 1000 permutation tests. Positive weights implied greater brain region volumes with higher behavioral scores, while negative weights indicated reduced volumes with higher behavioral scores.

## Results

### Demographics of the cohort and clinical characteristics

Initially, we enrolled 171 pediatric patients diagnosed with brainstem tumors and 849 typically developing children from 8 different sites. After excluding cases with poor MRI scanning (6 patients, 8 healthy children), those with tumors involving the cerebellum or thalamus (6 patients), and those with segmentation issues (10 patients), our final analysis comprised 147 patients, with a mean age of 8.5 years and 77 males. Among these patients, 114 had tumors located in the pontine region, 18 in the medulla oblongata, and 15 in the midbrain. We constructed a normative model using data from 849 typically developing children, with a mean age of 11.44 years old and 477 males (see Table 1). Fig. 3 shows the flow chart of the whole experiment.

**Figure 3.**
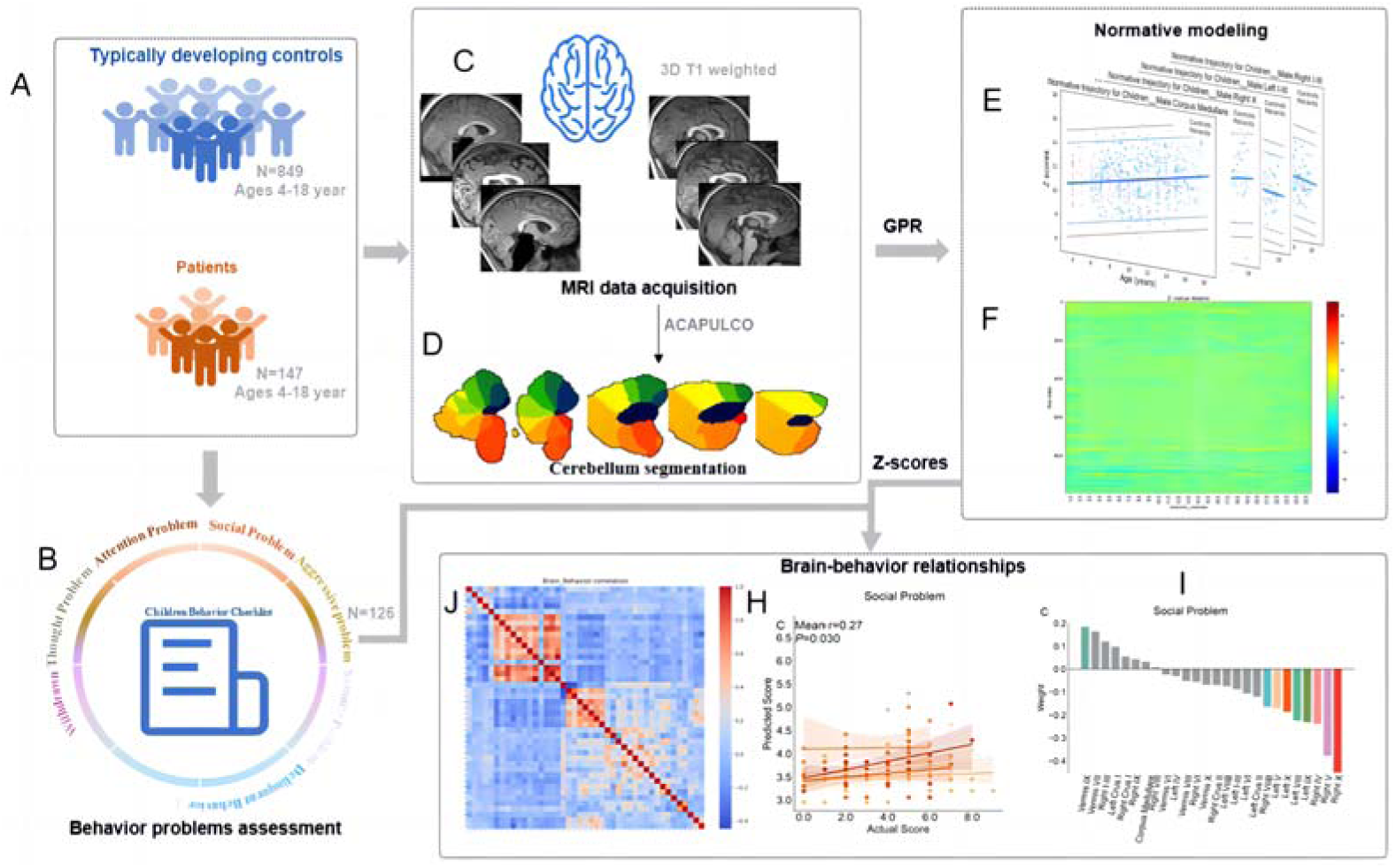
Data Analysis Flowchart. **(A)** Patient and Multi-site Healthy Children enrolled. **(B)** Patients’ behaviors were assessed in the neurosurgical clinic using the Child Behavior Checklist. **(C)** Structural MRI (T1) data acquisition and processing. Patients’ brainstem was excluded before segmentation. **(D)** All participants’ 28 cerebellar regions were obtained by Automatic Cerebellum Anatomical Parcellation using U-Net with Locally Constrained Optimization. **(E)** Normative trajectory in the GPR Model. In each model, a Gaussian model of the volume of each brain region was established with age and sex as covariates. **(F)** The Z-score profile of individual cerebellar regions’ volume. **(J)** Pearson correlation matrix of cerebellar regions and behavioral problems. **(H)** The deviations of cerebellar volume predict behaviors. (I) Feature weight maps for 26 cerebellar regions.

### Individual deviation of cerebellar volumes in patients

The normative trajectory derived from the GPR model for four cerebellar regions with the most extreme deviation demonstrated that patients’ abnormal deviation values primarily fell below the lower boundary of the general norm (Fig. 4A). The violin chart indicates that the majority of both patients and healthy children exhibited zero extreme positive deviations, with only 2.5% of subjects showing such deviations. In healthy children, tOC values were minimal and tightly clustered around zero. In contrast, patients exhibited a broader distribution of tOC values, with 95% of cases ranging between 3 and 10. Female patients exhibited a higher prevalence of extreme negative deviations compared to males (Fig. 4B). The stacked histogram suggested that patients primarily displayed positive deviations in the corpus medullare, while negative deviations were predominantly observed in Left I-III, Left V, Right V, Right I-III, Right X, Left X, Left IV, Vermis VI, Right IV, Vermis VII, and Right VIII (Fig. 4C).

**Figure 4.**
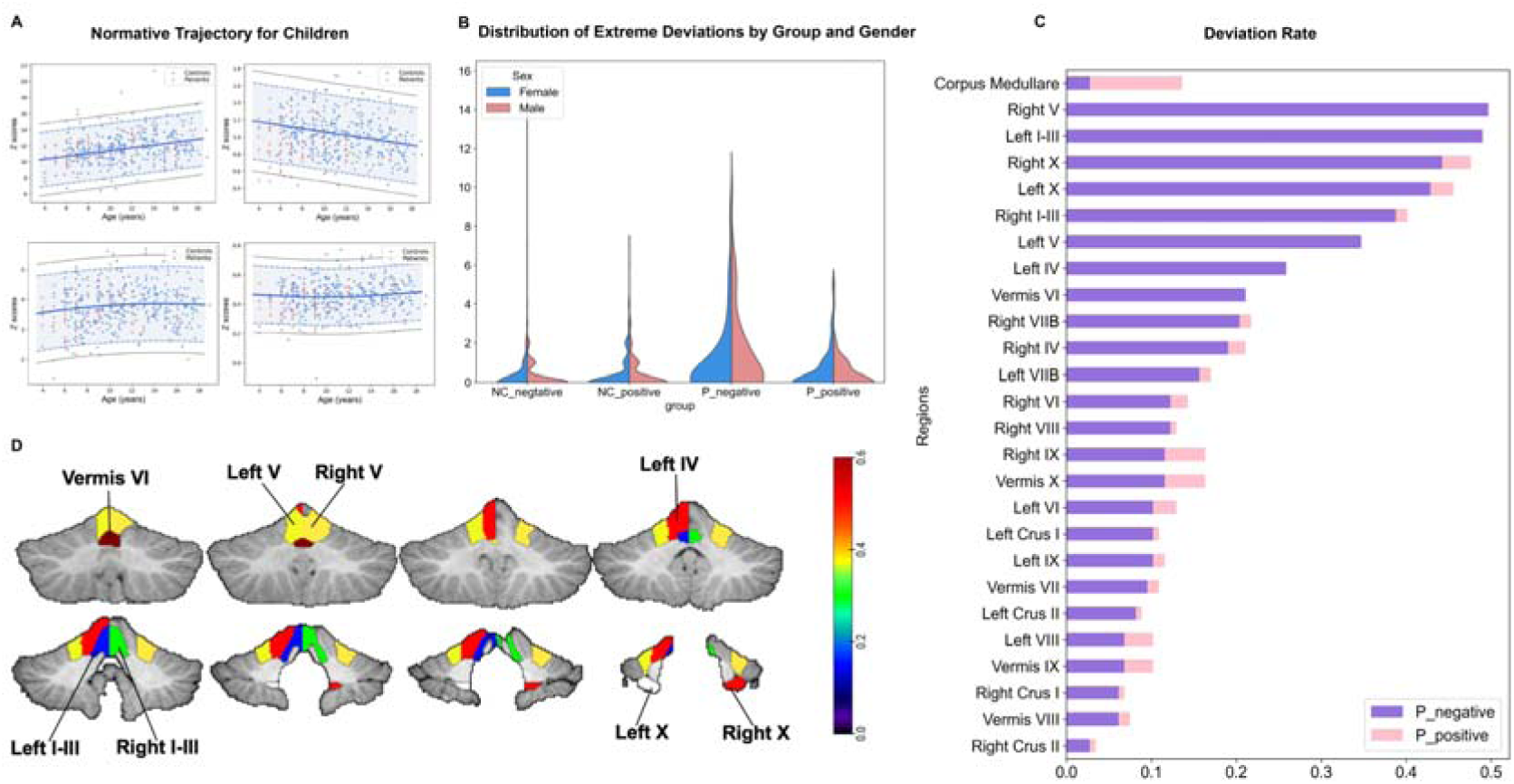
Normative model and the analysis of deviations. **(A)** The normative model was estimated using GPR. For the volume of 26 cerebellar lobes, the normative models were estimated. In each model, the volume of cerebellar regions was the response variable, and age and gender were covariates. These four subplots respectively show the normative trajectory for male patients and controls, with red dots representing patients and blue dots representing controls. **(B)** The violin plots display differences of positive and negative deviations in gender between patients and controls. **(C)** Stacking histograms of 26 cerebellar regions with positive and negative deviation rates in patients. The blue and red bars represent negative and positive deviation, respectively. **(D)** Illustration of extreme negative deviation of cerebellar regions (deviation rate >0.2).

### Outliers are associated with cognitive function

Our findings reveal a significant predictive relationship between the tOC and individuals’ behavior scores. Specifically, the analysis revealed a significant correlation between tOC and cognitive function, specifically with social problems (r = 0.31, p = 0.001) and withdrawal behaviors (r = 0.28, p = 0.001) as shown in Fig. 5A and 5B. Additionally, tOC was found to predict scores related to social problems (mean r = 0.27, p = 0.030) and withdrawal behaviors (mean r = 0.25, p = 0.034) as illustrated in Fig. 5C and 5D. These positive linear relationships indicate increased severity of social problems and withdrawal behaviors are associated with higher tOC values, which represent more severe cerebellar damage.

**Figure 5.**
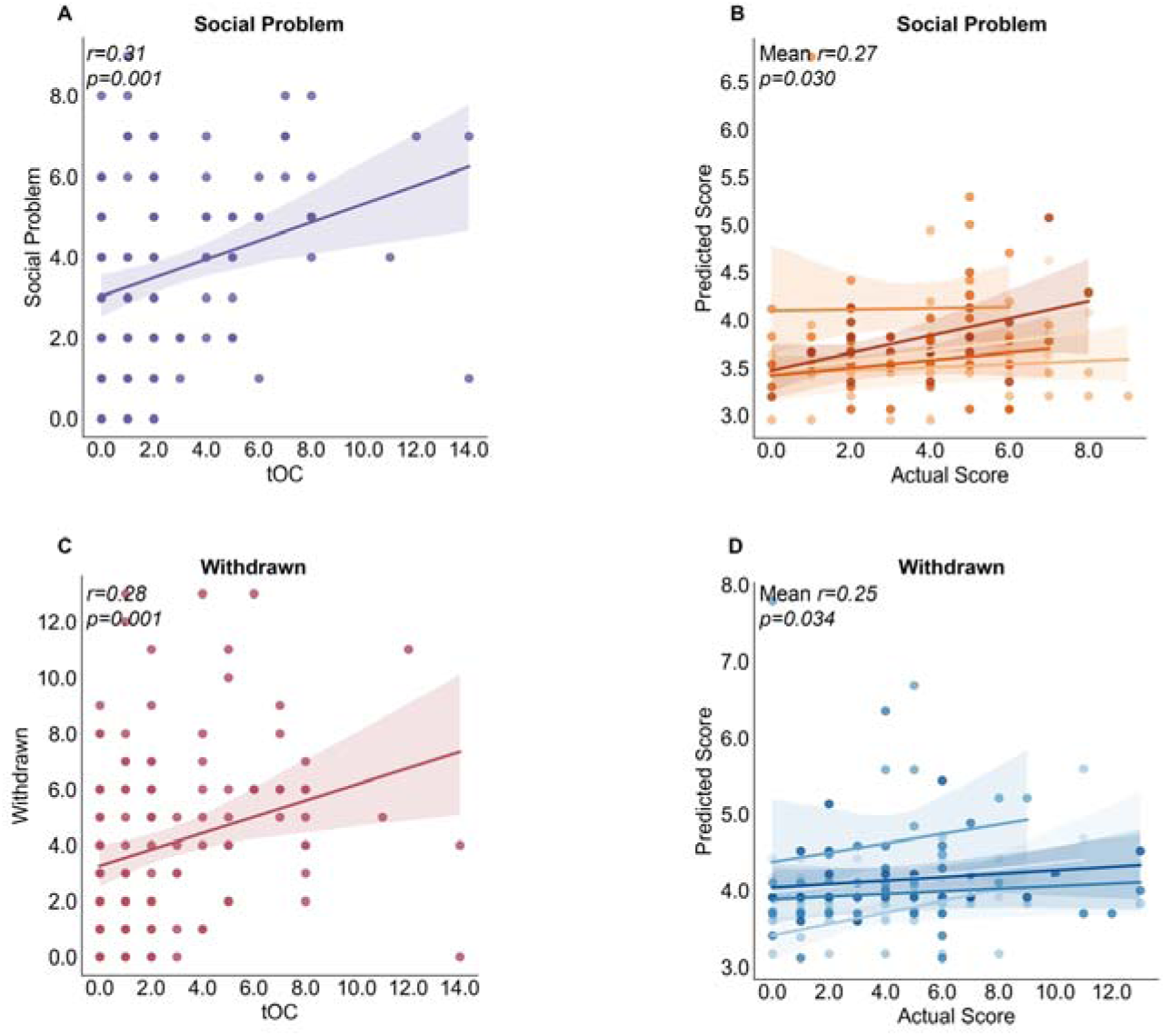
Behavioral prediction models using individualized tOC. The tOC predicts behaviors of glioma patients, including social problems and withdrawal behaviors. Scatter plots show the relationship between tOC and social problems (A), as well as withdrawal behaviors (B). tOC was also used to predict social problems (C) and withdrawal behaviors (D) in previously unseen individuals. Data points represent predicted scores based on five-fold cross-validation of independent test data, with p-values obtained through permutation testing.

### Predictive relationship between cerebellar volume deviations and behavior in glioma patients

Using the ridge regression model, we discovered a significant predictive association between cerebellar volume deviations and individuals’ behavior scores. Specifically, our analysis revealed that cerebellar volume deviations could predict scores associated with social problems (r = 0.27, p=0.007) (Fig. 6A), and withdrawal behaviors (r = 0.25, p = 0.015) (Fig. 6C).

**Figure 6.**
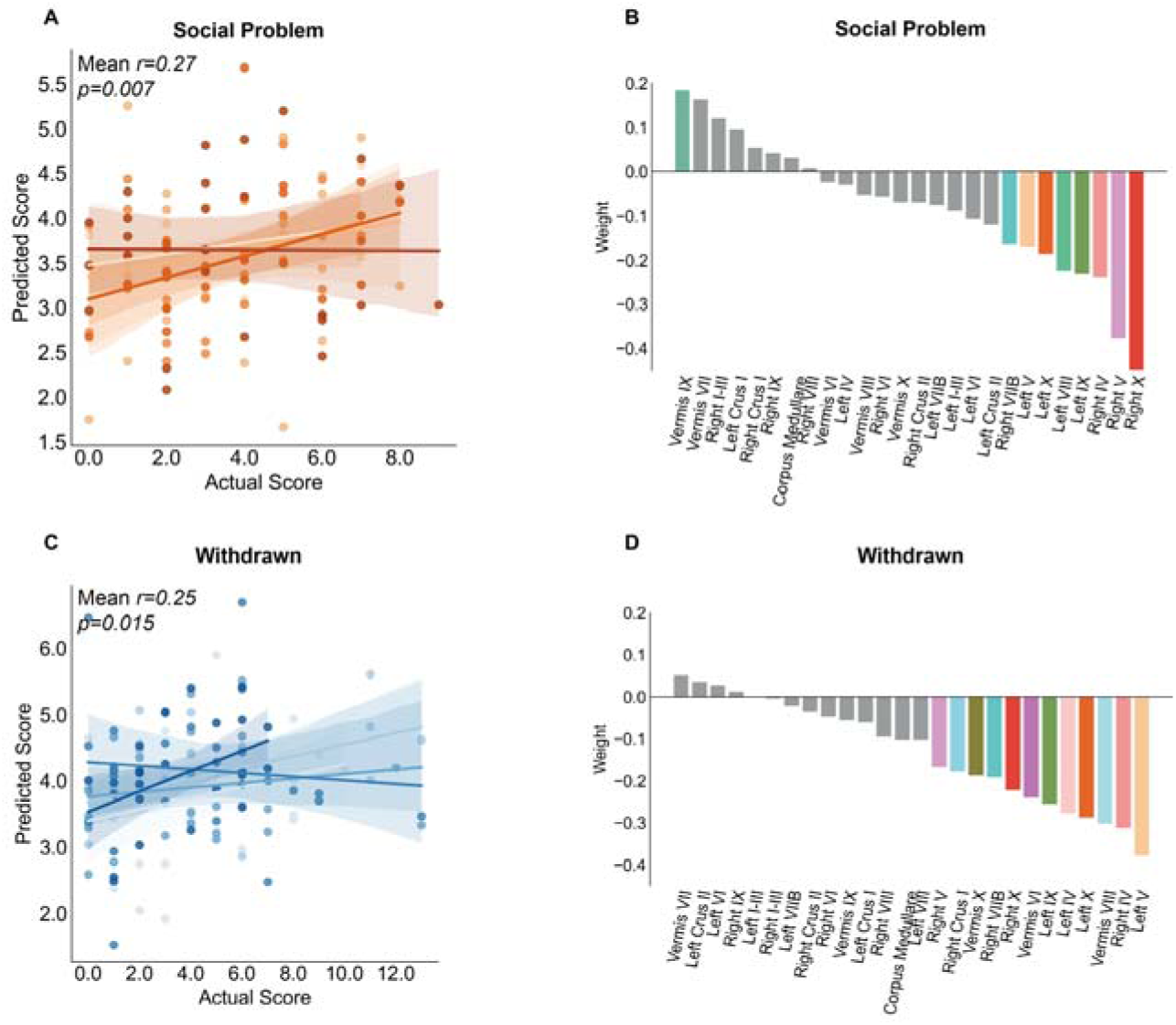
Behavioral prediction models of individual deviation of cerebellar volume. The deviations in cerebellar volume predict major behaviors observed in glioma patients, including social problems **(A)** and withdrawal behaviors **(B)**. Each data point on the graph represents the predicted scores (y-axis) of participants in a model trained on independent data using five-fold cross-validation. In each scatter plot, the five colors represent participants of the five subsets, respectively. The prediction accuracy of behavior is mean correction r between five folds. The p-value predicted for each behavior is significant. They are feature weight maps for 26 brain regions. The gray bar graph indicates that the p-value of the weight of this feature is not significant **(C-D)**. It indicated reduced cerebellum volume in Right X, Right V, Right IV, Left IX, Left VIII, Left X, Left V, and Right VIIB drove the prediction of social problem **(C)**. This revealed that reduced cerebellum volume in Left V, Right IV, Vermis VII, Left X, Left IV, Left IX, Vermis VI, Right X, Right VIIB, Vermis X, Right Crus I, and Right V drove the prediction of withdrawal behaviors **(D)**.

We observed that the region contribution weights predominantly displayed negative values, indicating reduced cerebellar volume in regions corresponding to more severe symptoms. For example, negative feature weights were evident in regions such as Right X, Right V, Right IV, Left IX, Left VIII, Left X, Left V, and Right VIIB for social problems (Fig. 6B), and in regions like Left V, Right IV, Vermis VIII, Left X, Left IV, Left IX, Vermis VI, Right X, Right VIIB, Vermis X, Right Crus I, and Right V for withdrawal behaviors (Fig. 6D). Furthermore, negative feature weights were consistent across both correlated behaviors in regions including Right X, Left X, Right V, Left V, Right IV, and Left IX.

## Discussion

This investigation revealed a general reduction in cerebellar gray matter among patients diagnosed with brainstem tumors, demonstrating a heterogeneous pattern of atrophy. Significant negative deviations were observed in Left I-V, right I-V, and Vermis IV-VII. Additionally, a negative correlation was identified between cerebellar lobe deviation and both withdrawal behavior and social problems. Moreover, the total outlier score emerged as an individualized indicator of cerebellar damage. In summary, the aforementioned findings yield the following insights: 1) brainstem tumors disrupt the normal neuroanatomical development of the cerebellum, 2) behavioral issues in children with brainstem tumors may be linked to abnormal cerebellar structures, 3) individual indicators of cerebellar lobular atrophy hold promise as predictors of behavioral outcomes. In the following sections, each of these findings will be further discussed.

### Cerebellar atrophy patterns in children with brainstem tumors

Severe atrophy predominantly affects the anterior lobes and vermis cerebelli, with milder impacts on the posterior and lateral lobes. This pattern likely arises primarily from compression due to the swollen brainstem. The current findings support clinical research highlighting the occurrence of secondary cerebellar atrophy induced by brainstem tumors. For example, previous studies reported significant cerebellar atrophy in patients with basal gangliocytoma involving the cerebellar peduncles^15^. Interestingly, cerebellar atrophy is common in patients with brainstem diseases. Recent research shows distinctive patterns of decreased cerebellar grey matter in patients with pontine infarctions, emphasizing lesion lateralization^27^.

The cerebellum undergoes rapid growth during the third trimester and continues to grow during adolescence with increased cellular processes and white matter proliferation. The volume of microstructures in the cerebellar peduncles increases until age 30^28^. Various factors, such as cerebellar hemorrhage, tumors, and indirect damages like diaschisis can lead to cerebellar volume loss. Studies on traumatic brain injury and animal models have shown cerebellar atrophy even without direct damage to the cerebellum^29^. The mechanism behind cerebellar atrophy can be attributed to three main reasons. Firstly, compression of the posterior fossa space by the swollen brainstem may limit normal neurodevelopment of cerebellum^30^. Secondly, disruptions in anatomical pathways can cause cerebellar atrophy. Impairment of pontocerebellar fibers can obstruct axonal transport between the cerebellum and brainstem, damaging the cerebellar tract and deforming cerebellar neurons ^15^. Thirdly, interruption of transsynaptic communication between pontine and cerebellar regions impacts fiber projection from cerebellar relay neurons to Purkinje cells in the cerebellar cortex, causing irreversible functional inactivation and long-term structural changes in the cerebellum^31^.

### Correlation between individual cerebellar deviation and behavior

Our study represents a pioneering effort to link secondary cerebellar changes caused by brainstem tumors to behavioral patterns in pediatric patients. We utilized tOC as a quantitative metric to assess cerebellar atrophy at the individual level. The tOC has shown high stability and reliability in predicting neuroanatomical heterogeneity among patients with Alzheimer’s disease^25^. We found that heightened cerebellar atrophy corresponds to increased withdrawal behavior and social problems in children. Using tOC as a metric for evaluating cerebellar health in individual patients could inform clinical decision-making. This discovery highlights the cerebellum’s role in regulating social functions and provides evidence of its significance as a core brain region^32^.

Furthermore, our study found that negative deviations in the volume of cerebellar subregions were significantly associated with worse social behaviors. Specifically, the volume of posterior cerebellar regions like lobules X, IX, and VIII, exhibited a significant negative correlation with social problems. These findings are consistent with previous studies, reinforcing the notion that the posterior cerebellar lobes contribute to social functions^33^. Moreover, the involvement of specific regions in the anterior cerebellar lobe, particularly lobules IV and V linked to the motor network, suggests that social functions are not limited to a single cerebellar area. This implies that social functions may be distributed across various areas of the cerebellum. For instance, functional MRI during social cognitive tasks has revealed activation in posterior cerebellar regions alongside the somatosensory motor network^32^. This connection highlights the complexity of cerebellar involvement in social functions. Furthermore, animal studies demonstrate that disrupted activity in anterior lobule IV/V leads to deficits in object recognition and social associative recognition tests, further supporting the cerebellum’s role in cognitive and social processing^34^.

A longstanding hypothesis in cerebellar research suggests that social dysfunction originates from damage in the vermis and posterior cerebellum^35^. This aligns with the dichotomous theory of cerebellar functions, which distinguishes between sensorimotor and cognitive-affective regions based on anatomical connectivity patterns. The spinal cord, brainstem, and sensorimotor-associated cortical regions prominently connect with the anterior cerebellar lobe and lobule VIII. Lesser connectivity extends to lobule VI and the interpositus nucleus^36^. However, the latest perspective suggests that the cerebello-thalamo-cortical (CTC) and cerebello-ponto-cortical (CPC) tracts establish connections between the posterior cerebellum, Crus I/II, and the social regions located in the frontal lobe^37^. Consequently, researchers hypothesize that injury along this pathway manifests as social deficits, with damage around the CTC and CPC tracts predicting social and emotional disorders^38^.

Moreover, it is important to note that the anterior and posterior cerebellar lobes, along with the vermis, show strong correlations with withdrawn behavior. The anterior cerebellar lobe contributes most significantly to this association. This suggests that withdrawal behaviors may arise from dysfunction in the brain’s social decision-making network^39^. The cerebellum and brainstem are integral components of the social decision-making network, contributing primarily through their regulation of somatic motor nuclei^40^. The present study suggests that the pathological withdrawal behaviors observed in patients may be attributed to severe atrophy of cerebellar motor nuclei. This finding corroborates numerous clinical observations indicating that children with lesions in the posterior fossa exhibit more complex and severe social problems, including withdrawal behaviors, attention deficit, irritability, pathological crying and laughing, and compression^41^. Importantly, extensive damage to the brainstem and cerebellar pedunculus results in more pronounced social dysfunction, with these behavioral disorders being profound and enduring^42^.

### Limitations of the study

This study is the first to investigate individual cerebellar changes in patients with brainstem tumors using automated segmentation techniques with the U-Net model. Unlike previous studies that relied on basic diameter measurements or subjective assessments, our approach provides precise quantification of cerebellar atrophy, allowing for larger-scale clinical research. By utilizing the normative model, our study mitigates clinical and biological heterogeneity, enhancing the robustness of our findings. A major contribution of our study is establishing a link between cerebellar damage from brainstem tumors and social behavior. While the impact of cerebellar injury on social functioning is known, neuroimaging evidence of cerebellar damage has been limited. Our study shows that damage to cerebellar regions around CPC and CTC tracts directly impairs social functions, expanding the focus beyond the cerebellar vermis. Additionally, the employment of ridge regression provides valuable insights for large-scale neuroimaging studies in clinical practice.

However, this study has some limitations. Firstly, the normative model only included age and gender-independent variables to estimate brain readouts. However, factors such as education and home environment may also affect the brain indices. Therefore, future, more comprehensive models should include these covariates to conduct a more comprehensive analysis. Secondly, the questionnaire utilized in this study primarily focused on assessing social problems. A more specialized tool may capture the nuanced aspects of these patients’ social functions. Thirdly, our findings reveal an association between cerebellar deformation and behavioral disturbance in pediatric patients. The nature of this relationship does not establish causality but rather implies a correlative linkage^43^.

## Conclusions

This study found significant individual heterogeneity regarding the impact of brainstem tumors on the cerebellum. Despite its prevalence, this aspect is often underemphasized due to the rarity of brainstem tumors. Our study presents a quantitative approach to assess this variability at the individual level. We analyzed spatial patterns of deviations in cerebellar region-of-interest (ROI) volumes. Individual neuroanatomical outlier profiles are linked to behavioral disorders. Notably, the count of outliers and volume of cerebellar lobules contribute to predicting social behavioral issues. These findings highlight the role of brainstem and cerebellar circuits in social behavior modulation. The robust correlation found between tumor-induced anatomical disruptions and cognitive functions advances tumor neuroscience research. Functional assessments before surgery are crucial for diagnosis, treatment, and preserving neural pathways during interventions.

### Author Contributions

Conceptualization, L.W.Z, L.Z.F.; Methodology, C.Y.C., H.Y.J., K.K.W., X.W.; Data acquisition, P.Z., M.X.Z., G.G.C, Y.Y.M., Y.X.T.; Formal Analysis, H.Y.J., K.K.W.; Writing - H.Y.J, K.K.W.; Writing - Review and Editing, All authors; Supervision, F.L.Z., P.Z., and L.W.Z.

## Acknowledgments

We are grateful to the patients and their families for making brains available for research. We also express our gratitude to Bianka Rumi for her thorough proofreading of the entire text.

## Ethical approval

All procedures performed in studies involving human participants were in accordance with the ethical standards of the institutional and/or national research committee and with the 1964 Helsinki declaration and its later amendments or comparable ethical standards.

## Data availability

Key resource table are available online and raw image data can be obtained upon reasonable request from the corresponding author.

