## Supplementary figures and images for "Individualized Cerebellar Damage Predicts the Behavioral Disorders in Children with Brainstem Tumors"

### Supplementary Figure 1

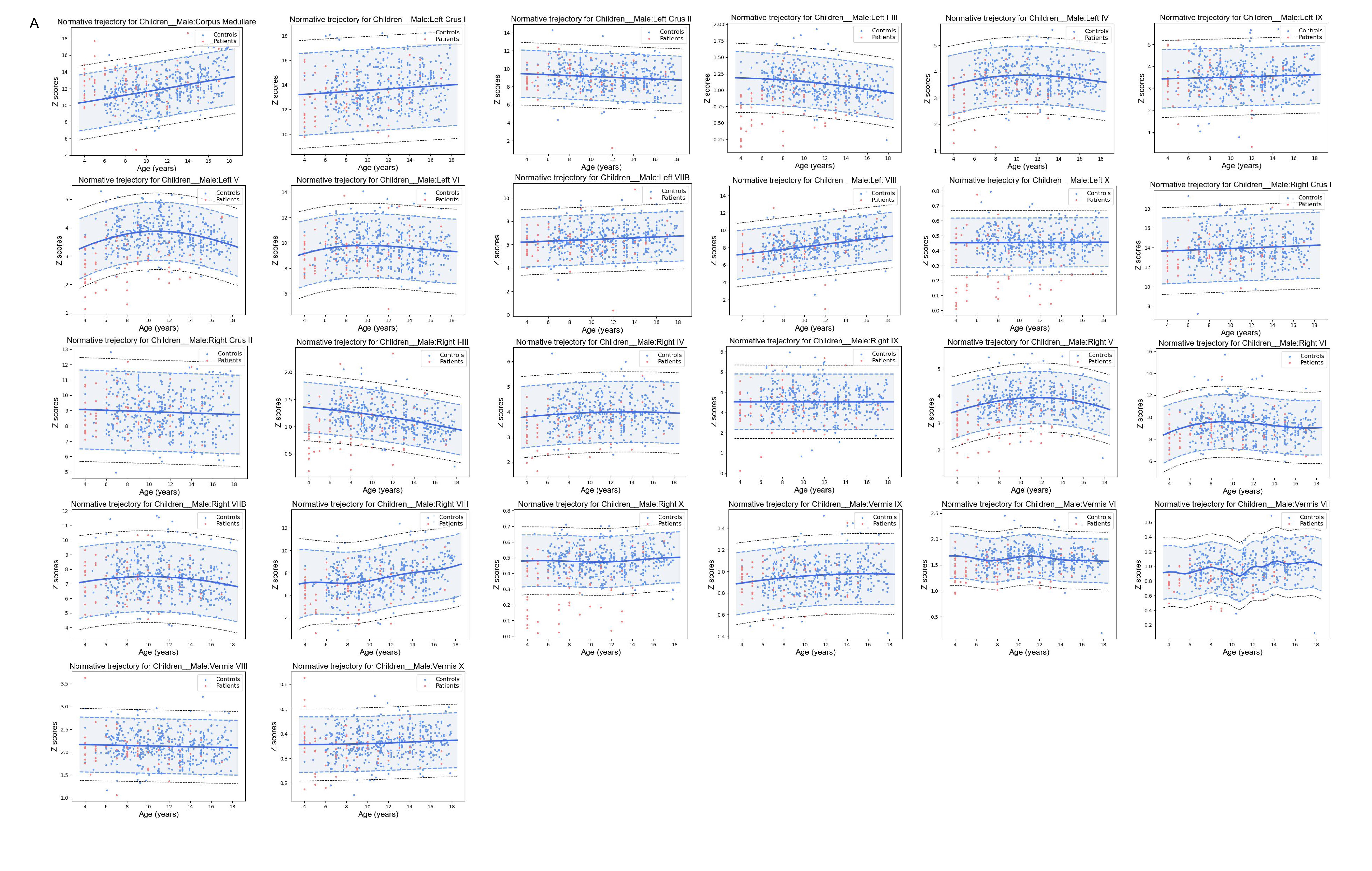
